# Opsonin-free, real-time imaging of *Cryptococcus neoformans* capsule during budding

**DOI:** 10.1101/313429

**Authors:** Hugo Costa Paes, Stefânia de Oliveira Frazão, Camila Pereira Rosa, Patrícia Albuquerque, Arturo Casadevall, Maria Sueli Soares Felipe, André Moraes Nicola

## Abstract

*Cryptococcus neoformans* is a unicellular fungal pathogen that causes meningoencephalitis, killing hundreds of thousands of patients each year. Its most distinctive characteristic is a polysaccharide capsule that envelops the whole cell. It is the major virulence attribute and the antigen for serologic diagnosis. We have developed a method for easy observation of the capsule and its growth dynamics using the cell-separation reagent Percoll and differential interference contrast (DIC) microscopy. Percoll suspension is far less disruptive of cell physiology than methods relying on antibody binding to the capsule, and measurements made with it are equivalent with India ink. Time-lapse microscopy observations using this method suggest that during budding, a dividing cell can regulate whether the capsule polysaccharide it produces is deposited on the capsule of the bud or on its own. This observation has important implications for our understanding of the *C. neoformans* capsule induction process during budding.

**List of abbreviations and acronyms:**

CSF: Cerebrospinal fluid
DIC: Differential interference microscopy
NA: Numerical aperture
CCD: Charge-coupled device
MM: Minimal medium
CIM: CO_2_-independent medium
MOPS: 3-Morpholinopropane-1-sulfonic acid
SD: Standard deviation

## Manuscript text

*Cryptococcus neoformans*, the most important agent of fungal meningitis, produces a thick polysaccharide capsule, one of its main virulence factors. This structure envelops the yeast cell, protects it from phagocytosis and has immunomodulatory properties that favor progression of the infection. The capsule is also the morphological signature of the *Cryptococcus* genus: the detection of yeast cells surrounded by a polysaccharide layer in cerebrospinal fluid (CSF) from patients is a standard procedure for diagnosis of cryptococcosis. For visualizing the capsule, the most common procedure is the India ink test: the CSF sample is mixed with a small amount of ink and a droplet of the resulting suspension is observed on a glass slide under a light microscope. The India ink particles form a dark background that reveals the light-permeant capsule around yeast cells by contrast (figure 1A). India ink staining is simple, cheap and quick, and a commonly used standard for researchers who wish to visualize the capsule.

**Figure 1.**
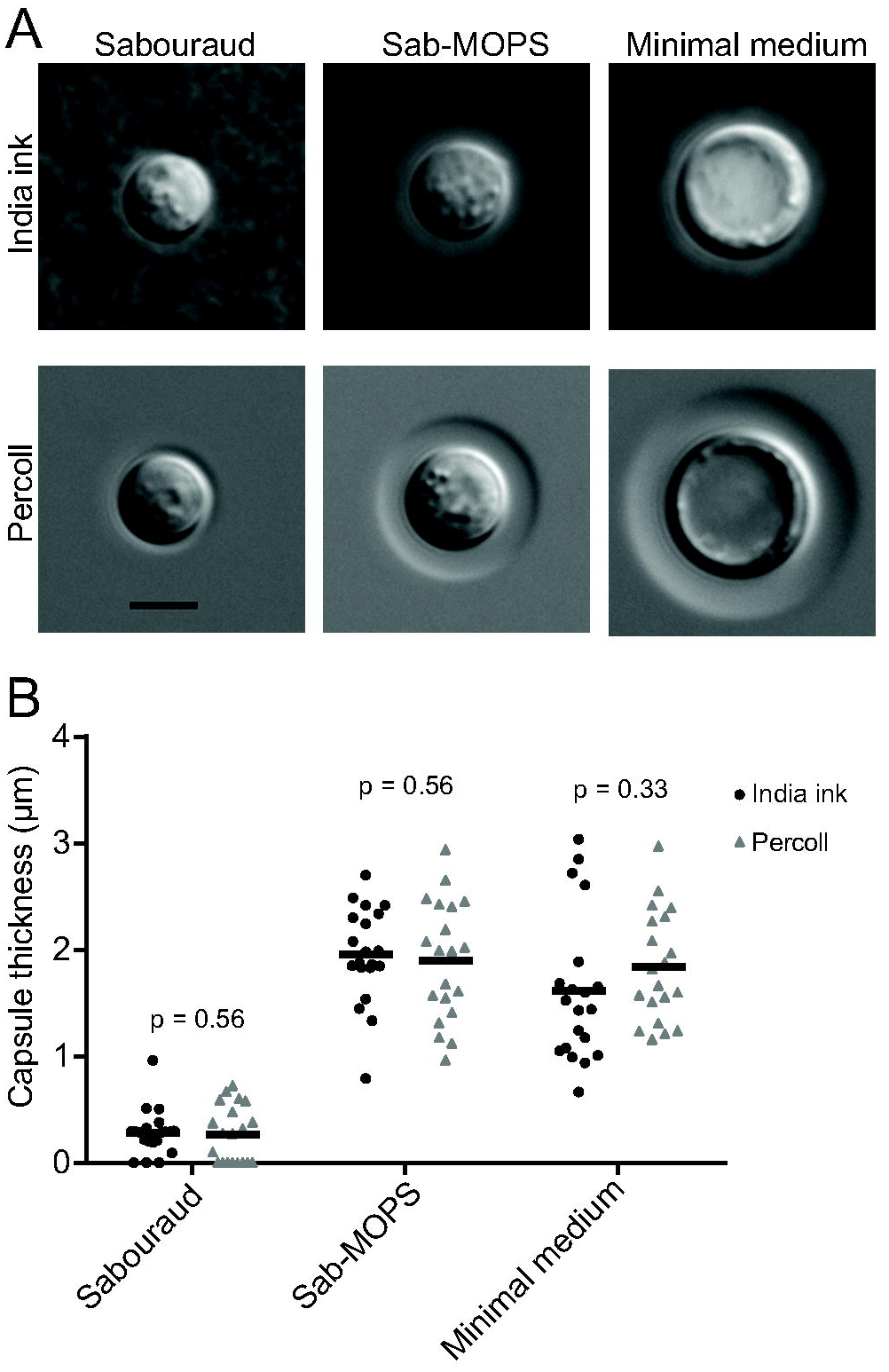
Comparison between DIC imaging with Percoll and India ink staining. *C. neoformans* cells were incubated in two different capsule-inducing conditions (MM and Sab-MOPS) and in non-inducing Sabouraud medium for 24 h prior to imaging with the method we describe and the standard India ink staining. Panel A shows representative images, whereas panel B shows that capsule thickness measurements yield similar results with both methods. The scale bar in A equals 5 μm. The lines in B represent the mean capsule thickness of 20 individual cells. The p-values shown in B were calculated by the Kolmogorov-Smirnov test comparing measurements with India ink and Percoll.

Despite its usefulness as a diagnostic technique, the India ink procedure has a major disadvantage for capsule research: commercial vendors add preservatives, such as thimerosal, to the ink preparation. This makes India ink formulations toxic for *C. neoformans* and only useful for static observations of the capsule. Consequently, India ink preparations cannot be used to document capsule formation in real time and, as a result, important aspects of the synthesis of this key virulence factor may escape observation. Some strategies have been used in the past to circumvent this limitation. *C. neoformans* capsules labelled with monoclonal antibodies^1^ or complement^2^ were incubated in capsule-inducing conditions and at specific time points during induction the probes were detected using fluorescence microscopy. More recently, the Quellung reaction observed on differential interference microscopy (DIC), which produces a change in refractive index when antibodies bind to the capsule^3,4^, was used to monitor capsule thickening in *C. neoformans* cells in real time^5,6^. These reports yielded important insights into the kinetics of capsule synthesis, but all suffered from an important caveat: because antibodies and complement proteins interact directly with capsule components, they may change its physical properties, especially at the reasonably high concentrations needed for the observations. Indeed, antibody binding increases rigidity of the capsule to a point that can actually hinder budding of daughter cells^7^ and it induces metabolic alterations in the fungal cell^8^. Hence, it is possible that these antibody effects affected the kinetics of capsule growth and there is a need for additional approaches to study capsule dynamics.

We hypothesized that a non-toxic suspension of particles with refractive index that is higher than that of the capsule would permit the observation of the capsule directly using DIC, as the particles would be excluded and create an optical path length gradient that generates contrast. We tested this hypothesis using Percoll, which consists of a 23% (w/w) aqueous suspension of colloidal silica particles (15-30 nm in diameter) coated with polyvinylpyrrolidone that is routinely used to form gradients for differential centrifugation of cells. As shown below, suspending *C. neoformans* cells in Percoll enabled us readily to observe the capsule as a well-defined halo around *C. neoformans* cells in high-resolution time-lapse DIC microscopy.

The microscopic observations were made on a Zeiss AxioObserver Z1 temperature-controlled inverted microscope equipped with 63X NA 1.4 oil immersion objective, DIC polarizers and prisms, motorized focus and an MRm cooled CCD camera (Carl Zeiss GmbH, Germany). The images shown below were collected using the Zeiss ZEN software and manipulated using Adobe Photoshop CS6, Adobe Illustrator CS6 and ImageJ. No non-linear modifications were made to the original images. *C. neoformans* cells of the H99 reference strain were incubated in either Sabouraud dextrose broth or one of three capsule-inducing media: 1) MM - minimal medium^9^ (29.4 mM KH_2_PO_4_, 10 mM MgSO_4_, 13 mM glycine, 3 μM thiamine, 15 mM dextrose) 2) CIM - CO_2_-independent medium^10^; Thermo Fisher Scientific, Waltham, MA, USA); 3) Sab-MOPS - Sabouraud broth diluted ten-fold with 50 mM 3-(N-Morpholino)propanesulfonic acid, pH 7.5^11^. All experiments were carried out at 37 °C. Proof of principle observations of *C. neoformans* incubated in Sabouraud broth, MM or Sab-MOPS for 24 h confirmed that the capsule was clearly visible (figure 1A) and that measurements made with India ink and Percoll produced equivalent results (figure 1B). Interestingly, we were also able to observe an increase in cell body diameter in cells incubated in MM, as reported previously^12^. In these experiments, we mixed *C. neoformans* cultures with equal volumes of India ink or Percoll and mounted the suspensions on slides covered with 0.170 mm coverslips. To test whether Percoll was toxic to *C. neoformans*, we set up parallel cultures with and without Percoll for 84 h and estimated final cell densities on a hemocytometer: these were 5.2 x 10^7^ cells/ml (SD: 3.9 x 10^6^ cells/ml) and 6.4 x 10^7^ cells/ml (SD: 1.3 x 10^7^ cells/ml), respectively (p=0.2004). The results established that suspension in Percoll was not toxic for growth. We observed that the capsule edge was much sharper in Percoll suspension than in India ink suspension. Although the exact mechanism for this difference is not known, we suggest two explanations that are not mutually exclusive. First, India Ink is a preparation of particles with heterogeneous sizes and some of the smaller particles may be able to penetrate the domain of the capsule, especially given that outer layers are less dense^2^. Second, the capsule is a highly hydrated and hydrophilic structure that may not interact well with the polyvinylpyrrolidone thus creating a sharp exclusion zone. These encouraging results led us to set up experiments that would allow us to document capsule formation in real time by time-lapse microscopy.

For time-lapse imaging, we observed cells in capsule-inducing media supplemented with Percoll in a POCmini-2 cultivation system (PeCon GmbH, Germany), a closed chamber bounded by two 0.170 mm coverslips. In some experiments we observed that a large proportion of cells in a 50% Percoll suspension would float due to being less dense than the medium. Thus, we enriched the suspension for denser cells by resuspending cells into a 50% Percoll suspension in water (v/v) and centrifuging at 2000 *g* for five minutes. The pellet containing cells that did not float was then counted and suspended at low density (approximately 1000 – 2000 cells per mL) in CIM or Sab-MOPS and supplemented with 50% (v/v) Percoll. MM is not suitable for time-lapse experiments because the mixture turned into a gel after prolonged incubation. Approximately 800 μL of the resulting suspension were then added to the chamber, which was incubated in the microscope at 37 °C to allow collection of images every five to fifteen minutes for one to two days. The time-lapse images show that the capsule growth is more pronounced in Sab-MOPS, although it was readily observable in both capsule induction conditions (figures 2A and 2B and supplemental videos 1-3). We measured the capsule size for several cells and found that the capsule growth follows sigmoidal kinetics with different rates for different cells (figure 2C), as previously observed using measurements based on the Quellung effect^5,6^.

**Figure 2.**
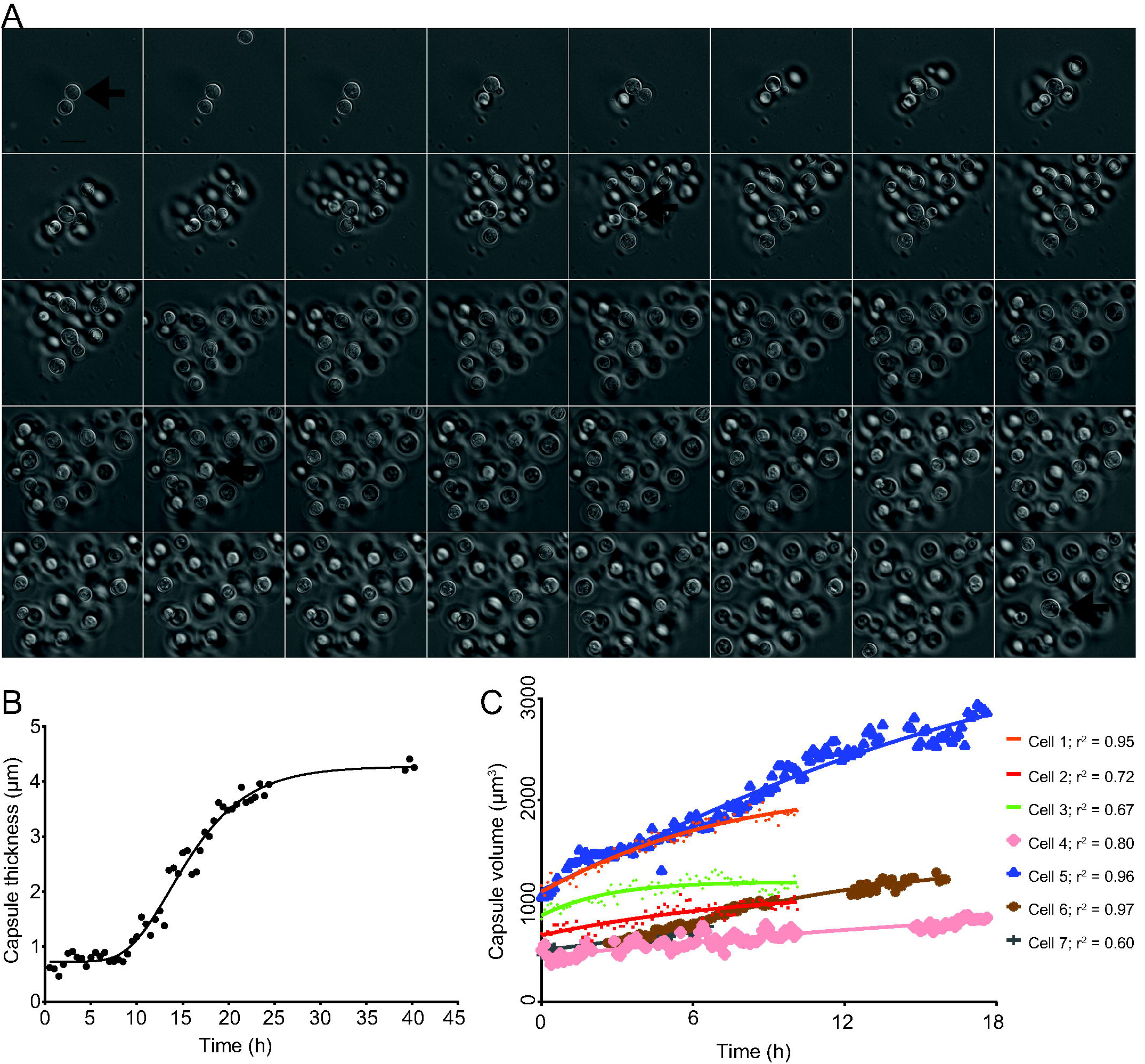
Time-lapse imaging of *C. neoformans* capsule growth in inducing medium (Sab-MOPS) *C. neoformans* cells were incubated in Sab-MOPS supplemented with 50% Percoll and imaged every 10 minutes by DIC microscopy. The images that were used to make this figure were compiled in supplemental video 1. Panel A shows some of the images, with a 60-minute interval between each image. The arrows point to the same cell at different time points; this cell was used to make figures 2B and 3B. Panel B shows the capsule thickness measured every 30 minutes and the line represents an asymmetric sigmoidal curve that was fit to the data (r^2^ = 0.98). In panel C, capsule volume was calculated at several time points for seven different cells using cell body and capsule diameter measurements from micrographs as described in the text. Curves were fitted to the values using the Gompertz growth curve model in Graphpad Prism 6 (adjusted r^2^ values shown). The curves generated for each dataset were compared to test for variation in the slope (*k*) and the preferred model, with a probability greater than 99.99%, was found to be that *k* is different for each curve. Gaps in some curves correspond to periods in the time series when the cells drifted out of focus, making it impossible to perform measurements.

In contrast to the studies using antibodies, however, we were clearly able to observe the capsule in nascent buds and follow its growth in daughter cells. The capsule measurements shown in figure 3A and plotted in figure 3B are from a cell that went through six budding events throughout the observation period. This cell is indicated in figure 2A by black arrows. The first three buds emerged during the phase in which the mother cell capsule was not growing; all three had capsules that were thicker than that of the mother cell and each one had a thicker capsule than the previous bud. The later buds, in contrast, emerged as the mother cell capsule was becoming thicker; their capsules were less thick than those of the mother cell and the previous buds. In contrast to this one cell, we observed several others whose buds had a thin capsule and no apparent pattern of capsule redistribution from their mother cell. These observations suggest that the destination of new capsular material synthesized by the mother cell – its own capsule or that of a bud – may not be stochastic. In a zebrafish model of cryptococcal infection, *C. neoformans* cells with enlarged capsules effectively inhibited macrophage phagocytosis and caused more severe disease^13^. Capsular polysaccharide protects the cell from phagocytosis and free radical fluxes^14^. Thus, whether capsular material is destined to increase the size of the mother cell or the bud could define which of these cells escapes the immune response inside the host.

**Figure 3.**
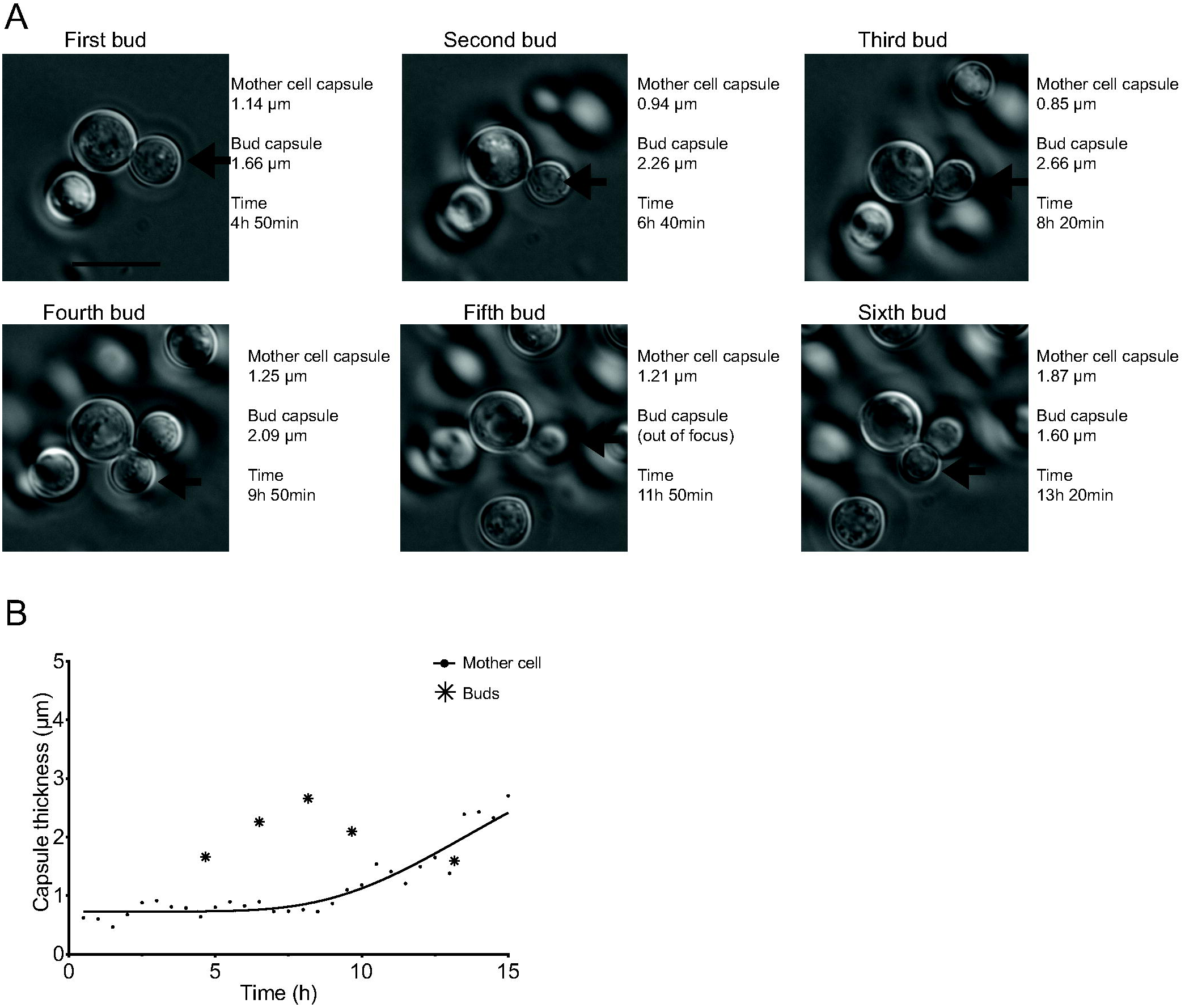
Comparison between the capsule thicknesses of one mother cell and it multiple buds. Panel A shows the same mother cell indicated by arrows on figure 2A, which appears supplemental video 1, but at the specific time points in which each of six successive buds (arrows) can first be seen to be completely separated from the mother cell. The actual images, capsule measurements and how much time had elapsed after the cells were transferred to capsule inducing media are indicated in the figure. In panel B, the bud capsule sizes are plotted into a zoomed-in version of the curve shown in figure 2B, which shows capsule growth kinetics for the mother cell.

Another interesting pattern we observed in several (but as seen in figure 3 not all) budding cells on Sab-MOPS is of mother cells keeping their capsules at a constant thickness while budding and increasing it afterwards (figure 4). When we compare our data including buds to those of García-Rodas *et al.^6^*, at first glance they seem to be in agreement: while the mother cell is budding actively, and presumably spending only brief intervals in G1/S, its own capsule does not become thicker. Our data would seem to suggest that this is less because of an intrinsic impairment of capsule growth during G2/M, as suggested by the authors of that study, and more because a mechanism exists whereby capsule polysaccharide might be destined for the capsule of the daughter cell.

**Figure 4.**
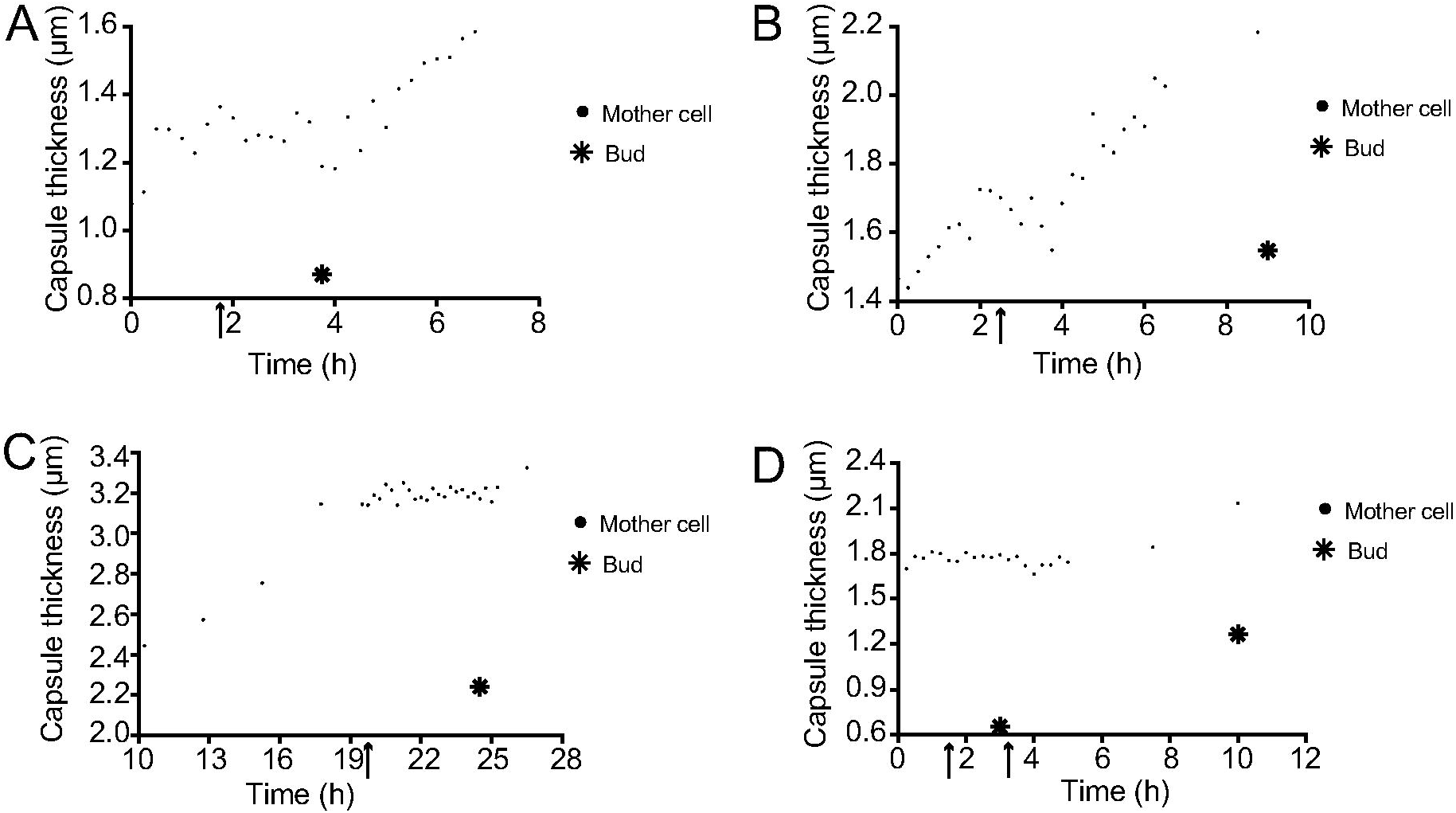
Capsule thicknesses of four mother cells and their buds. Panels A-D show the capsule thickness of four mother cells and their buds. The arrows point to the first time point in which the bud can first be seen emerging in the cell wall of its mother cell, whereas the star indicates the bud’s capsule thickness at the first time point in which its cell wall has clearly separated from that of its mother cell.

Comparing India ink staining and the method we describe here, we found that the former uses a cheaper reagent and requires a much simpler microscope for observation. However, India ink is not readily available everywhere and the quality of the reagent for capsule visualization varies widely among manufacturers. Despite its higher price, Percoll resulted in consistently successful observation of capsule by different researchers in our lab. More importantly, it allowed real-time observation of the capsule in live cells in the absence of exogenous protein binding to – and potentially interfering with – the capsule. Thus, while the need for a DIC-equipped microscope precludes its use in clinical diagnosis of cryptococcosis, we propose that Percoll could become the contrast agent of choice for research laboratories that have access to appropriate equipment.

In summary, we describe a new method for visualizing the capsule and studying capsular growth kinetics based on suspending cells in media with a very different refractive index than the capsule. Comparison of the insights obtained with this method relative to those learned with the Quellung method confirms that different cells grow at different rates. Perhaps most importantly, the Percoll method has provided new insights on the distribution of polysaccharide during budding that have important implications for our views on bud survival and antibody function.

## Acknowledgments

The authors would like to thank the Brazilian funding agencies Capes, CNPq and FAP-DF for research funding and scholarships.

## Captions for the supplementary videos

**Supplementary video 1 – Capsule growth in Sab-MOPS**

*C. neoformans* cells were re-suspended in capsule-inducing medium (Sab-MOPS) supplemented with 50% Percoll and imaged every 10 min for 40 h. The time-lapse images were then cropped and compiled into a video. Still images from this video are shown in figure 2A, and several details of one of the cells in this video (along with its buds) are shown in figures 2B, 3A and 3B.

**Supplementary video 2 – Capsule growth in CIM**

*C. neoformans* cells were re-suspended in capsule-inducing medium (CIM) supplemented with 50% Percoll and imaged every 5 min for a total period of 23 h.

**Supplementary video 3 – Capsule growth in CIM**

*C. neoformans* cells were re-suspended in capsule-inducing medium (CIM) supplemented with 50% Percoll and imaged every 5 min for a total period of 23 h.

